# Sequential Activation of Senescence and Apoptosis by ONG41008 Eliminated the Human Pancreatic Ductile Adenocarcinoma in nude mice and xenografted human lung cancer

**DOI:** 10.1101/2025.06.03.657005

**Authors:** Byung-Soo Youn, Kyung-Jin Kim, YunKyung Kim

**Affiliations:** Osteoneurogen, Inc.; Dae-Gu Kyung-Bug Institute for Science and Technology (DGIST); College of Pharmacy and Won Kwang University

**Keywords:** Senescence, Senescence-Coupled Apoptosis (SCA), Polyphenols, Anti-cancer drug, Lung Adenocarcinoma (NSCLCs), Pancreatic Ductal Adenocarcinoma (PDAC)

## Abstract

Current therapeutic limitations associated with pancreatic ductal adenocarcinoma (PDAC) are still evident. We have found that ONG41008, the first human synthetic polyphenol, is able to eradicate PDAC in a nude mouse model. ONG41008 is known to potently attenuate human TGF-β production, inducing both anti-fibrotic capacity and anticancer potential through senescence-coupled apoptosis (SCA). These interesting observations closely paralleled ONG41008’s ability to inhibit PDACs, and as expected, ONG41008 alone was able to actually ablate engrafted PDACs in nude mice. Interestingly, the anti-PD 1 developed by MSD did not appear to act on immune activation of exhausted Cd8+ T cells in the tumor microenvironment (TME), but primarily on engrafted PDAC blood vessels. These observations support the latest report from MSD. A549 cells known to be a well appreciated human lung adenocarcinoma cell line were treated with ONG41008. Consistent with the prior observations ONG41008 notably induced SCA. and when A549 cells were xenografted into mice daunting tumor formations were noticed. Upon stimulation with ONG41008 via direct intramural injection of these tumors’ tumor volumes were rapidly regressed.

Taken together, we believe that ONG41008 could be a multifaceted anti-cancer agent not only against PDAC but also against other aggressive cancers such as lung cancer.

## Introduction

Polyphenols are characterized by the presence of the conserved Chromone-Scaffold (CS) exclusively found in the plant kingdom (1). More than 10,000 kinds of polyphenols have been discovered. Multiple biological functions involving anti-inflammation have been attributed to polyphenols. Thus, polyphenols could be the next-generation therapeutic drug candidates for many human intractable diseases (2).

Drawing a rational implication of immunotherapies in real cancer modalities is yet to be done. More through scrutinization of the hitherto immunotherapies in a natural context of agonism and antagonism must be explored. We here show that ONG41008 alone would be able to regress and defragment the PDAC masses in nude mice via its multifaceted functions involving anti-fibrosis, anti-cancer (senescence-coupled apoptosis) (SCA), and anti-inflammation (3-5). Having been much in line with these discoveries, we believe that polyphenols would be able to open a new venue whereby most intractable cancers could be cured. In fact, A549, human lung adenocarcinoma cell line, is prone to apoptosis by ONG41008, and A549-xenograft tumors were remarkably attenuated by ONG41008.

## Results and Discussions

### ONG41008 is the first human synthetic polyphenol exhibiting multiple key biological manifestations in human intractable diseases

To demonstrate whether ONG41008 is an efficacious anti-cancer drug we decided to closely examine the chemical compositions of ONG41008. As seen in Figure 1A, ONG41008 harbors a typical Chromosome-Scaffold (CS) as a major chemical constituent which is linked to a benzyl methoxy group. We had found the methoxy located at C6, the 6th position of CS played a pivotal role in all the biological manifestations of ONG41008. While Quercetin and Fisetin were explored to be Senolytic compounds ONG41008 was a multifaceted functional polyphenol exhibiting anti-inflammation, anti-fibrogenesis, and anti-cancer. Pyocyanidin C1 was the latest senolytic drug presumably being a trimer. Seeing whether the trimeric structure is essential for its resistance to onset of aging remains to be intriguing. Apigenin is known to be a potent anti-inflammatory compound, and we showed that apigenin was able to mount inactivation of innate immunity of LPS-stimulated macrophages (data not shown). And Genistein has been well appreciated as a general tyrosine kinase inhibitor, suggesting that polyphenols existing in the plant king dome are a group of multifaceted functional compounds healing for many human intractable diseases.

**Figure 1.**
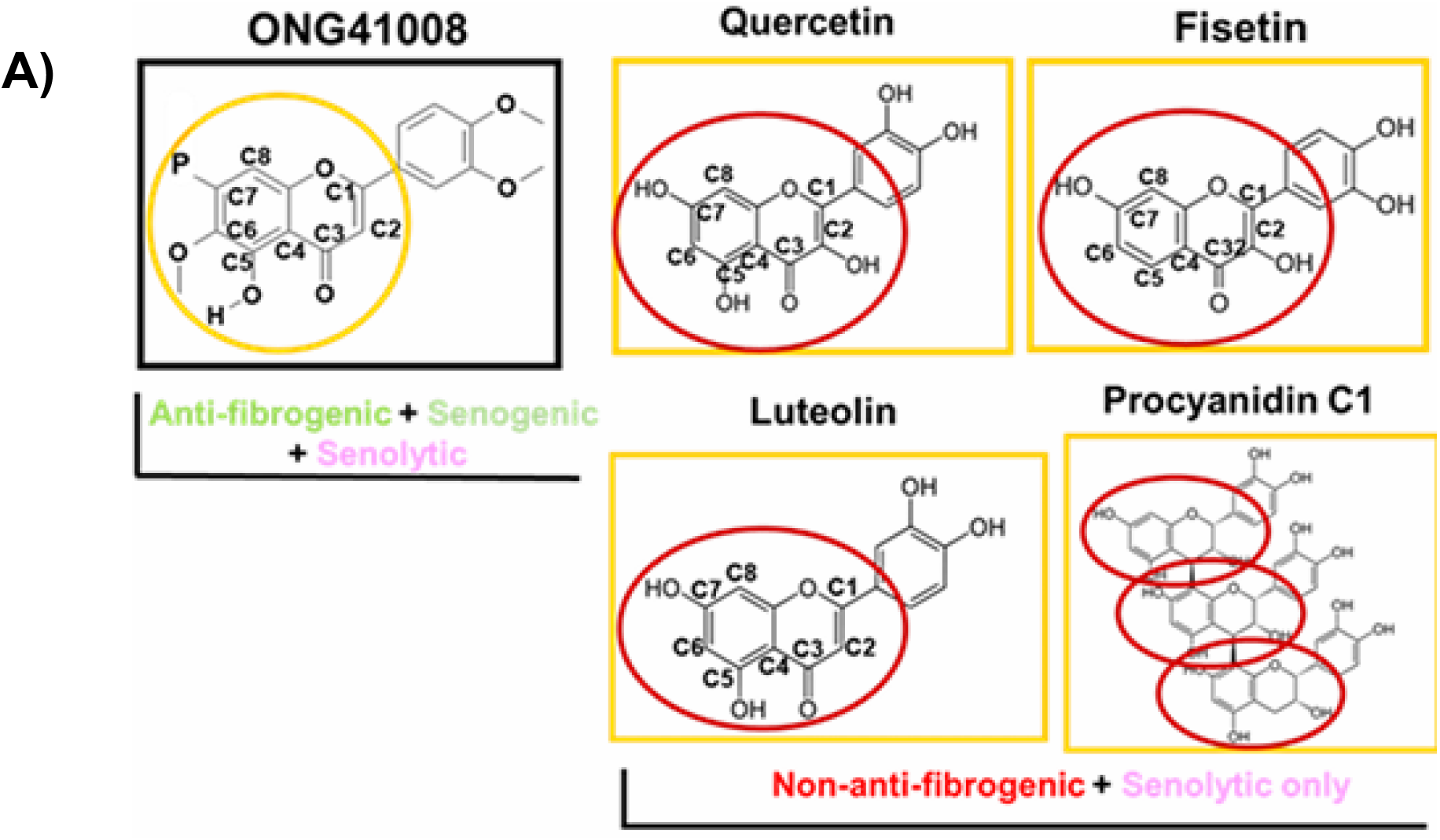
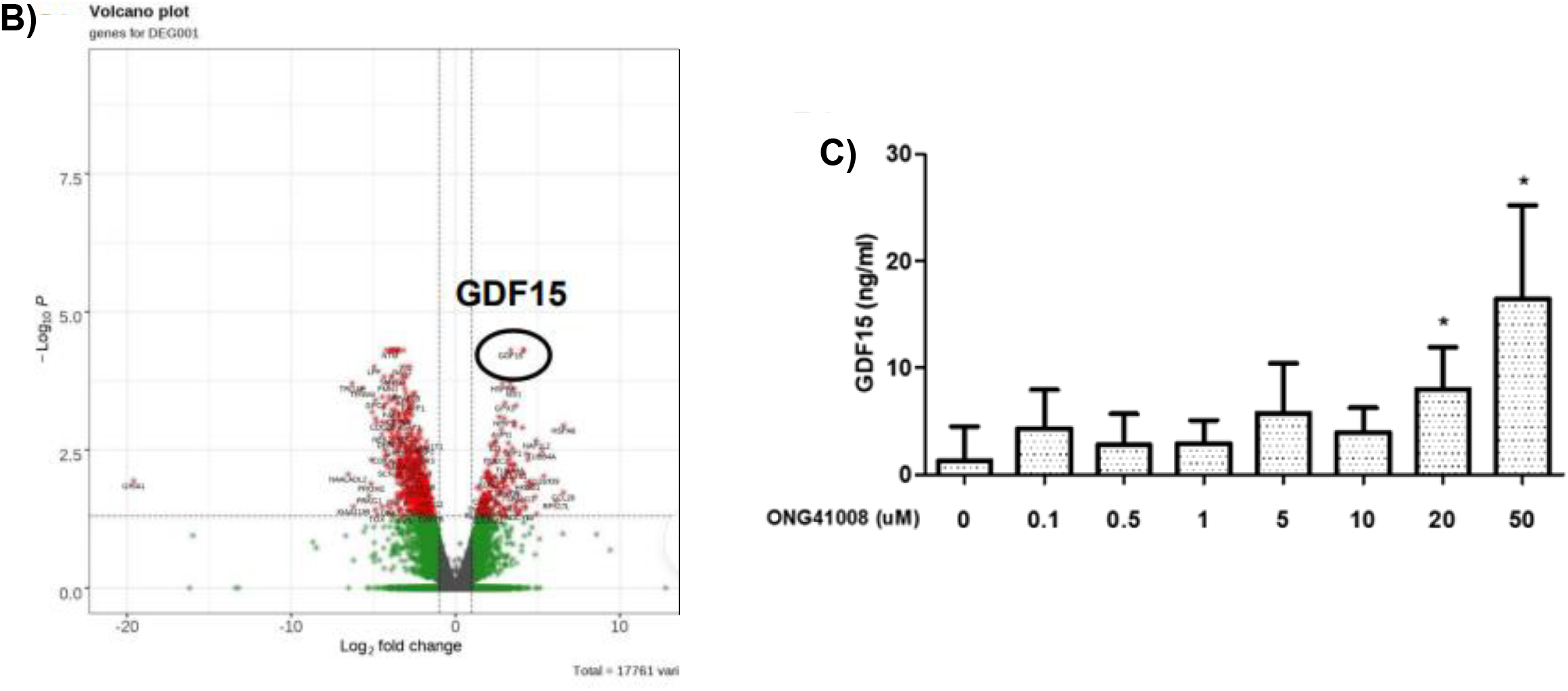
ONG41008 exhibits a typified polyphenol structure and shows a little diversification impacting on its multifaceted biological features. A) ONG41008 is anti-fibrogenic, senogenic and senolytic. B) Both quercetin and fisetin are senolytic. C) Apigenin is a potent inflammatory drug and genistein is a general tyrosine kinase inhibitor.

### ONG41008 is a potent inducer of senescence and apoptosis in A549, a human lung adenocarcinoma

We previously showed ONG41008 induced SCA in various human cancer cells (1). The SCA led to Mcl1 induction in PANC1, a human malignant PDAC cell line (1). Mcl1 induction should be related to two showrooms of Mcl1 which have been known to be responsible for apoptosis (4). A rapid onset of NAD synthesis presumably occurring in mitochondria in A549. An expression profiling before and after ONG41008 treatment was performed. Surprisingly, it occurred to us that ONG41008 bifurcated the incoming signaling into FAS receptor signaling and p16 activation that seemed to be mediated by induction of GDF15. Its biological meaning remains uncertain at the present time. To explore the induction of SCA in A549, the cells were stimulated with ONG41008, and induction of senescence and multinucleation of the stimulated A549 cells were observed in H2AX staining. As shown in Figure 2A, ONG41008 was able to rapidly induce H2AX expression as well as to enhance multinucleation. Furthermore, xenograft-transplantations of A549 cells in nude mice followed by direct intratumoral injection of ONG41008 were remarkably regressed tumor sizes and tumor volumes. We asked a question whether induction of Mcl1 was also required for the apoptosis of ONG41008-treated A549. However, differential recruitment of different Bcl2 family members seemed to be able to integrate cell death signals depending on the cell origins.

**Figure 2.**
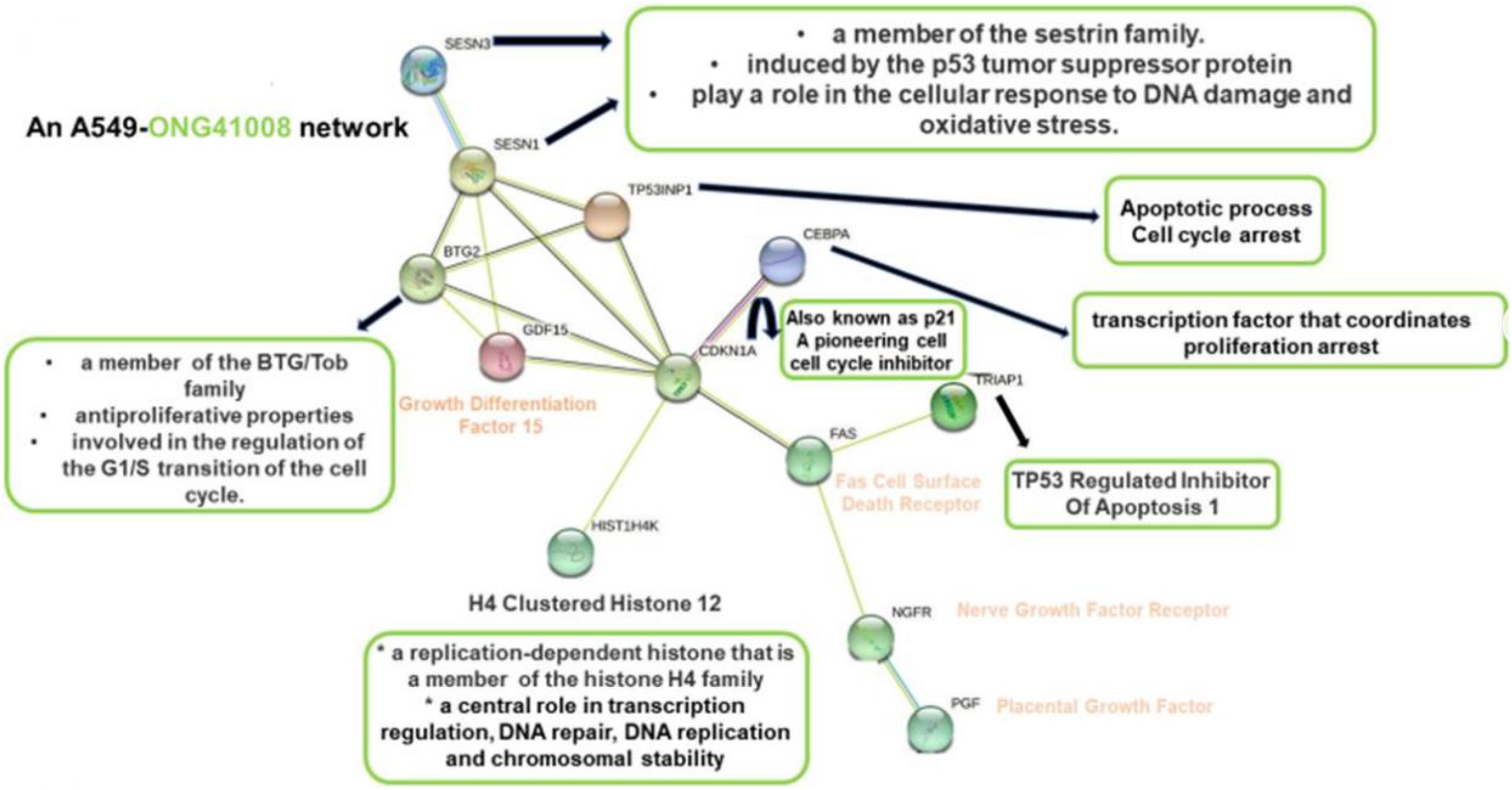
A) A549 cells were stimulated with 10uM ONG41008 for 72 hrs along with DMSO. F-actins and a senescence-inducible chromatin called H2AX were double-stained and visualized. B) Xenograft transplantation of A549 cells and direct administration of increasing concentrations up to 100 mpk of ONG41008 into established xenografted tumors.

**Figure 3.**
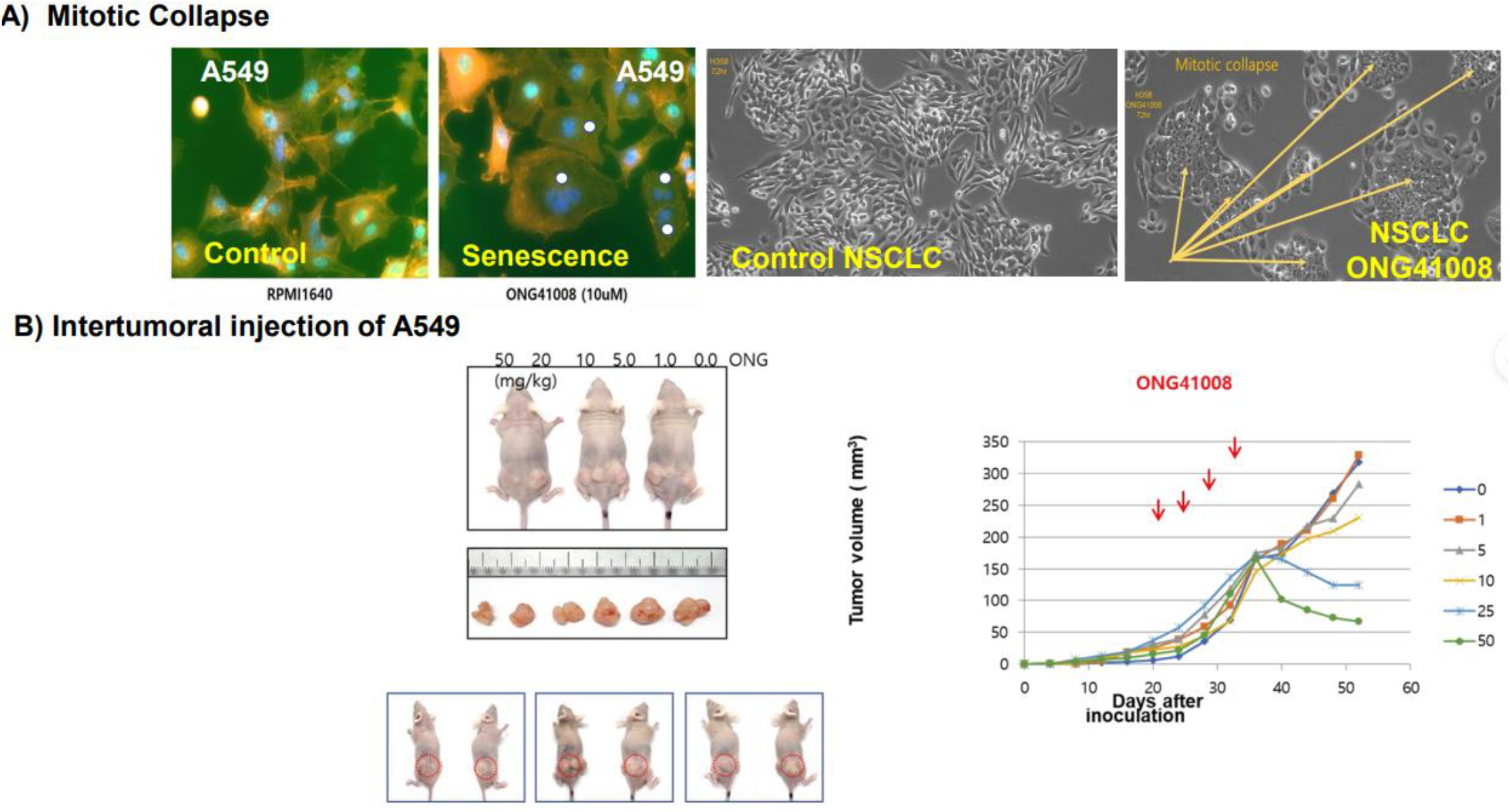
RNA-Seq analysis of ONG41008-treated A549 cells. Expression profiling and protein interactomes were scrutinized.

Taken together, ONG41008 can rapidly attenuate human lung cancer formation in xenograft mouse models.

### ONG41008 defragmented the PDAC-engrafted cancer mass independently of anti-PD1 in nude mice

ONG41008 seemed to be a self-contained anti-cancer drug due to its multiple biological advantages over human major intractable diseases. We simply established cancer elimination exploring set by using PDAC in nude mice models. As shown in Figure 4, as compared to control mice, ONG41008 alone substantially defragmented the established PDAC via anti-fibrosis after three to four weeks, and the entire cancer masses were dramatically reduced when time went by. Interestingly, anti-PD1, an MSD human monoclonal antibody, was also substantially able to regress cancer masses but yet to be eliminated, suggesting anti-PD1 seemed to act on the established cancer vasculatures possibly not on impaired T cell tumor immunities in TME. This interesting feature was well appreciated by an MSD report ().

**Figure 4.**
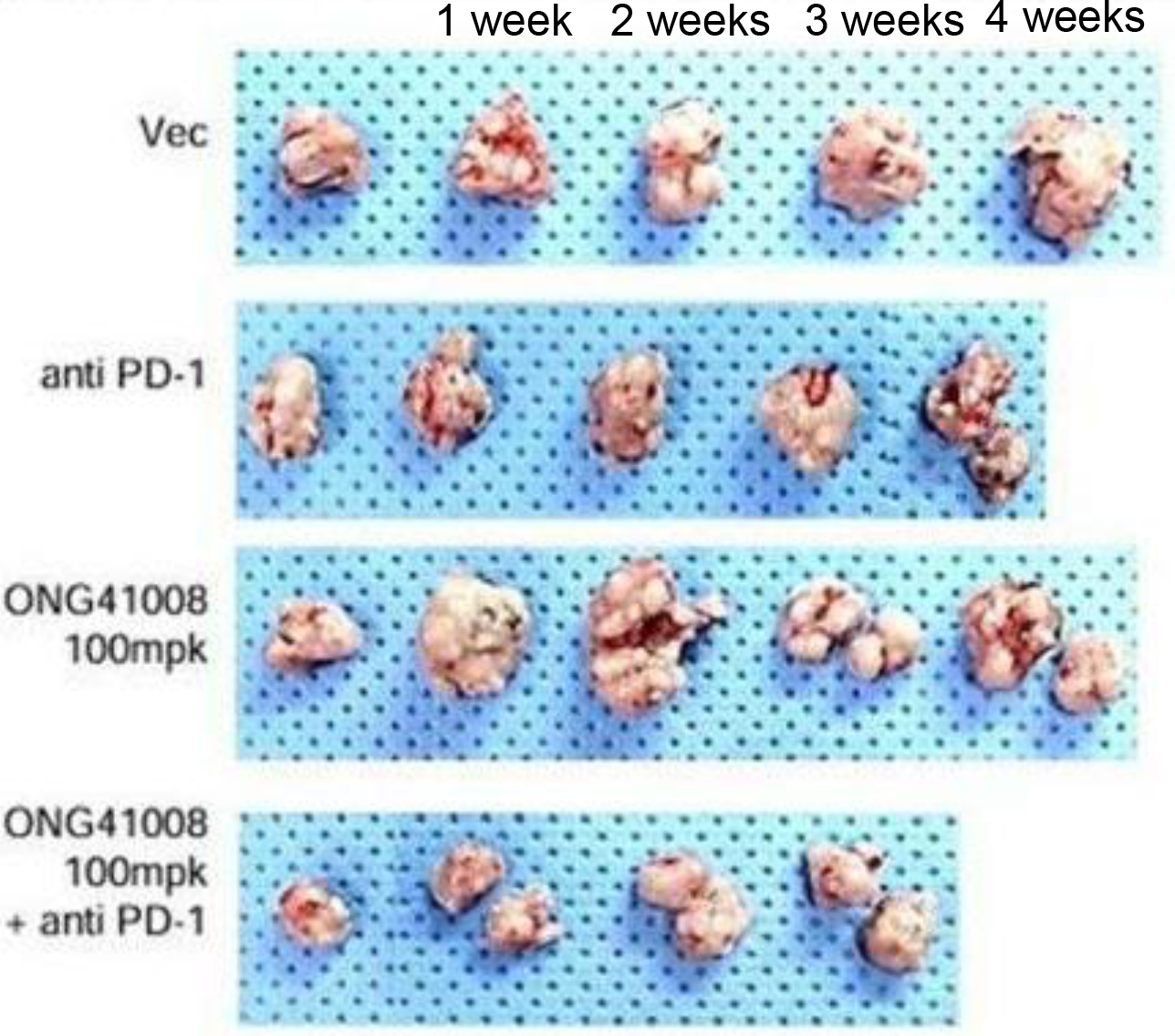
Elimination of human malignant PDAC into nude mice by ONG41008 Human primary aggressive PDAC was engrafted into nude mice and subjected to control, anti-PDA (100 ug), ONG41008 (100 mpk), and ONG41008 + anti-PD1. Cancer defragmentation and tumor vasculature were monitored.

Combined ONG41008 with anti-PD 1, a synergistic defragmentation followed by substantial reduction in tumor sizes of the PDAC-established tumor occurred but seemed unnecessary upon an extrapolation of the experiments up to 2 months. Once again, this data emphasized the importance of tumor vasculature and senescence-coupled apoptosis (SCA) in eliminating engrafted PDAC into immune-compromised mice. And, for the treatment of medically emergent PDAC patients could be treated with the combination of ONG41008 and anti-PD1, an MSD monoclonal antibody.

## Acknowledgement

A contractual agreement between Osteoneurogen and the Korea University Industry Collaboration was established for the investigation of ONG41008 and human lung cancer. A contractual agreement between Osteoneurogen and the Korea National Cancer Research Institute was made for the investigation of ONG41008 and PDAC. Fidelity Science Group (FSG) has been established under Osteoneurogen governance. Its voluntary members at the present time are Drs. B-S YOUN, KJ KIM, KS PARK, B-S KWON, JB KIM, J-S KIM, and Y-S KIM.

## Materials and Methods

### 1. Cell Culture and Reagents

The A549 cell line was purchased from the Korean Cell Line Bank (Seoul, Korea) and cultured in RPMI supplemented with 10% FBS and 1% P/S (Welgene, Seoul, Korea). Chemically synthesized ONG41008 was obtained from Syngene International Ltd. (Bangalore, India), dissolved at a stock concentration of 50 mM in DMSO, and stored in aliquots at −20 °C. DMSO was used as a control. PDAC cells were isolated from malignant human PDAC patients and were cultured in RPMI supplemented with 10% FBS and 1% P/S (Welgene, Seoul, Korea).

### 2. Immunocytochemistry

Cells were fixed using 4% paraformaldehyde, permeabilized with 0.4% TritonX100, blocked with 1% BSA, and incubated with rhodamine phalloidin (Thermo Fisher, Waltham, MA, USA), anti-p53 (Cell Signaling Technology, Beverly, MA, USA), p21(Abcam, Cambridge, UK), p16-INK4A (Proteintech, Rosemont, IL, USA) for 4 h at room temperature. After washing, cells were incubated with an Alexa Fluor 488 (Abcam, Cambridge, UK)-conjugated secondary antibody. Images were analyzed using EVOS M7000 (Invitrogen, Waltham, MA, U

### 3. Xenograft of A549 and syngeneic transplantation into nude mice

One million cancer cells were grown and transplanted into the corresponding mice. These mice were treated with control and ONG41008 (100mpk) via oral administration. For immunotherapy, 100ug anti-PD1 per mice was administered via IV. Tumor volume and masses were assessed.

### 4. RNA-seq, Differential Gene Expression, and Interactome Analyses

Processed reads were mapped to the Mus musculus reference genome (Ensembl 77) using Tophat and Cufflink with default parameters. Differential analysis was performed using Cuffdiff using default parameters. Then, FPKM values from Cuffdiff were normalized and quantitated using the R Package tag count comparison (TCC) to determine statistical significance (e.g., p-values) and differential expression (e.g., fold-changes). Gene expression values were plotted in various ways (i.e., scatter, MA, and volcano plots) using fold-change values and an R-script developed in-house. The protein interaction transfer procedure was performed using the STRING database with the differentially expressed genes. A 60 Gb sequence was generated, and 10,020 transcripts were read and compared. The highest confidence interaction score (0.9) was applied from the Mus musculus reference genome, and information regarding interactions was obtained based on text mining, experiments, and databases (http://www.string-db.org/ accessed on 6 December 2021).

